# Bacterial colonizers of *Nematostella vectensis* are initially selected by the host before interactions between bacteria determine further succession

**DOI:** 10.1101/2022.12.13.520252

**Authors:** H Domin, J Zimmermann, J Taubenheim, G Fuentes Reyes, L Saueressig, D Prasse, M Höppner, RA Schmitz, U Hentschel, C Kaleta, S Fraune

## Abstract

The microbiota of multicellular organisms undergoes considerable changes during development but the general mechanisms that control community assembly and succession are poorly understood. Here, we use bacterial recolonization experiments in *Nematostella vectensis* as a model to understand general mechanisms determining bacterial establishment and succession. We compared the dynamic establishment of the microbiome on the germfree host and on inert silica. Following the dynamic reconstruction of microbial communities on both substrates, we show that the initial colonization events are strongly influenced by the host but not by the tube, while the subsequent bacteria-bacteria interactions are the main cause of bacterial succession. Interestingly, the recolonization pattern on adult hosts resembles the ontogenetic colonization succession. This process occurs independently of the bacterial composition of the inoculum and can be followed at the level of individual bacteria, suggesting that priority effects are neglectable for early colonization events in *Nematostella*. To identify potential metabolic traits associated with initial colonization success and potential metabolic interactions among bacteria associated with bacterial succession, we reconstructed the metabolic networks of bacterial colonizers based on their genomes. These analyses revealed that bacterial metabolic capabilities reflect the recolonization pattern, and the degradation of chitin might be a selection factor during early colonization of the animal. Concurrently, transcriptomic analyses revealed that *Nematostella* possesses two chitin synthase genes, one of which is upregulated during early recolonization. Our results show that early colonization events are strongly controlled by the host while subsequent colonization depends on metabolic bacteria-bacteria interactions largely independent of host development.

## Introduction

All multicellular organisms live in association with microbes. These microbes can have a variety of effects and functions in metabolism (1), immunity (2), pathogen resistance (3), development (4) and behavior of their macroscopic host (5). The growing understanding of the effects of the microbiome on its host raises the questions of how microbial communities assemble, how they resist perturbation and how they function in the context of the host.

In host-microbe research, the complexity of the microbiome poses one of the biggest challenges. To facilitate questions about how beneficial communities assemble, several ecological models were applied to a microbial scale, such as niche theory and neutral theory (6), null models (7) or Vellend’s understanding of community assembly (8). Nemergut et al. updated Vellend’s framework on community assembly for microbial communities, explaining assembly processes through only four processes: selection, diversification, dispersal and drift, while also accounting for the differences between macrobial and microbial community assembly (8,9). While dispersal and drift are considered to be more stochastic and therefore mainly dictated by chance, selection and diversification are considered to be more deterministic and therefore mainly dictated by bacterial, environmental, or host factors that exert selection or diversification pressure.

Recently, *Nematostella vectensis* became popular as a model to understand host-microbe interactions (10). *Nematostella* is a cnidarian sea anemone belonging to the Anthozoans, and although cnidarians belong to the early-branching metazoans, *Nematostella* exhibits a surprisingly large genetic complexity, possessing most signaling pathways for development and immunity important in bilaterian animals (11,12). *Nematostella* readily undergoes its complete life cycle under laboratory conditions and its whole bacterial community composition was characterized over the course of its development, as well as its virome (13,14). Thereby, the microbiome of *Nematostella* changes with developmental age, shows spatial structuring along the body column, and exhibits a diurnal pattern (14–17). In doing so, it shows strong resistance to community overgrowth by one member (18). Recently, high microbial plasticity in response to environmental changes has been functionally linked to thermal adaptation in *Nematostella* (19).

Here, we aim to understand the fundamental principles underlying the establishment and succession of complex microbial consortia on host tissue. Our results show that the recolonization dynamics recapitulate ontogenetic colonization pattern of *Nematostella*, regardless of the initial composition of the inocula. Thereby, single members of the microbiome can be divided into early- and late-colonizing bacteria, which are defined by their appearance during recolonization. Early colonization correlated with a high abundance of polysaccharide degradation pathways, especially for potentially host-provided chitin. In agreement, transcriptomics analysis showed an increased expression of host chitin synthase genes. In contrast, late-appearing bacteria were increasingly capable of oxidizing compounds such as nitrite and sulfide which earlier colonizers potentially released by nitrate and sulfate reduction. Thus, we highlight the successive nature of bacterial colonization of the host *Nematostella* and suggest a role for host-microbe interactions via chitin as a driver of early colonization events and bacteria-bacteria interactions as a driver of later colonization events.

## Materials and Methods

### Animal culture

The adult animals of the laboratory culture were F1 offspring of CH2XCH6 individuals collected from the Rhode River in Maryland, United States (20,21). Animals were kept under constant, artificial conditions without substrate or light. For *Nematostella* Medium (NM), Red Sea Salt was diluted in Millipore H_2_O and adjusted to 18°C and 16‰ salinity. Feeding occurred 2-3 times a week with first instar nauplius larvae of *Artemia salina* (Ocean Nutrition Micro Artemia Cysts 430–3500 g, Coralsands, Wiesbaden, Germany). Primary polyps were fed with homogenized larvae until they were big enough to feed on whole larvae. Spawning was induced adapted after Genikhovich et al. 2009 by shifting the temperature to 25°C and exposure to light for 10 hours (22). Fertilization was performed *in vitro* in petri dishes by transferring the egg packages into NM containing sperm. Fertilization of the eggs was performed within one hour of release of the egg package from the mother.

### Antibiotic treatment

Antibiotic treatment was adapted after Domin & Gutiérrez et al. 2018 (18). Sterility was confirmed firstly via plating of homogenized polyps on marine broth (MB) plates. Absence of CFUs was interpreted as sterile. Secondly, sterility was checked via a PCR with primers specific for V1-V2 region of the bacterial 16S rRNA gene (27F and 338R). Although a slight band could be observed, no recovery of bacteria over the course of the experiment could be observed in subsequent PCRs and plating on MB plates, attributing the slight PCR band to dead bacterial matter.

### Recolonization

For the recolonization experiments of live polyps, the protocol for conventionalized recolonized H*ydra* polyps was modified (3). The germfree adult polyps were recolonized with the microbiota of three different developmental stages, respectively. For the four time points (2, 7, 14, and 28 days post recolonization), four germfree polyps were pooled in one vessel. Experiments were conducted with five independent replicates. For recolonization with adult stages, one adult polyp per one germfree polyp was homogenized (4 homogenized polyps/ 50 mL NM), for early and juvenile stages approximately 0.1 mL of animals per adult polyp were homogenized (0.4 mL homogenized animals/ 50 mL NM). Early stages were 6 days old, juvenile stages 54 days old. After 24 hours, the medium was exchanged to remove tissue debris and non-associated bacteria. After another 24 hours, samples for the first time point (2dpr) were collected. For each sample, one polyp was used. After washing the polyp three times, it got either homogenized in NM for gDNA extraction or frozen in liquid nitrogen for RNA extraction. For 16S rRNA-sequencing an extraction with the DNeasy Blood & Tissue Kit (Qiagen) was performed, for RNA-sequencing the RNA was extracted with the RNeasy Plant Mini Kit (Qiagen). If the animals were homogenized in NM, 1/100 of the homogenized animal were plated on marine broth plates prior to gDNA extraction and the plates were incubated for at least 2 days at 18°C to count CFUs. Due to extraction difficulties, the experiments for gDNA extraction and RNA extraction were performed separately.

For the recolonization of silicone tubes, hollow silicone tubes with an inner diameter of 3 mm, an outer diameter of 5 mm and a wall thickness of 1 mm were cut into 1 cm long pieces. Tubes were recolonized and sampled exactly like the adult polyps but with 10% MB in NM. For sampling, tubes were washed three times and bisected longitudinally. One half was used for gDNA extraction and 16S rRNA-sequencing, the other half was used for biofilm quantification with crystal violet. For this, the tubes were incubated in 1 mL of 0.1% crystal violet solution for 15 minutes. Afterwards, tubes were washed three times with water before the tubes were dried overnight. Then crystal violet was washed off the tubes with 500 μL of 95% ethanol for 15 minutes with slight agitation. Absorbance was measured at 550 nm.

### DNA extraction and 16S rRNA-sequencing

Prior to gDNA extraction, the animals were washed three times with 500 μL sterile NM and frozen without liquid at −20°C until extraction. The gDNA was extracted with the DNeasy Blood & Tissue Kit (Qiagen, Hilden, Germany) as described in the manufacturer’s protocol. DNA was eluted in 50 μL elution buffer. The eluate was frozen at −20°C until sequencing. Sequencing was conducted as described in (18). The raw data are deposited at the Sequence Read Archive (SRA) and available under the project ID PRJNA902551.

### 16S rRNA sequences processing

Filtering and taxonomic analysis were conducted according to the qiime2 pipeline (23,24). Sequence quality filtering was performed via DADA2 and taxonomic analysis via the q2-feature-classifier plugin for qiime2 with the Greengenes 13_8 97% OTU data set as reference (25–27). Further downstream analysis was conducted using the R package phyloseq (28) and plots were generated with the R package ggplot2 (29). The statistical tests adonis and anosim were calculated with the R package vegan (30). Because qiime2 creates the abundance table according to exact sequence variants (ESVs) and not operational taxonomic units (OTUs) on a specific identity percentage anymore, we manually clustered the ESVs into OTUS with 97% identity with cd-hit-est (31,32) for the metabolic pathway analysis. The output sequences were called clusters instead of ESV or OTU.

### Quantification of total bacterial abundance

In order to quantify the relative bacterial abundance in comparison to host tissue, we performed quantitative real time PCR with the 27F/338R bacterial primers, and primers for the elongation factor 1alpha gene (F GTAGGCCGTGTTGAGACTG, R CACGCTTGATATCCTTCACAG) of *Nematostella*. The expression levels were calculated according to the ΔΔCT method (33). We used the GoTaq qPCR Master Mix (Promega) with MicroAmp 0.2 mL optical strips (Applied Biosystems) and a QuantStudio 3 qPCR system (Applied Biosystems).

### RNA extraction and sequencing

Prior to RNA extraction, polyps were washed three times in sterile NM. After pipetting off as much liquid as possible, polyps were immediately frozen in liquid nitrogen and stored at −80°C until extraction. Total RNA was extracted with the RNeasy Plant Mini Kit (Qiagen) according to the manufacturer’s protocol. RNA was eluted in 30 μL RNase-free water that got reapplied on the column’s membrane and eluted again. RNA quality was checked via application on an agarose gel and measured on a Qubit. RNA libraries were constructed using the TruSeq stranded mRNA (incl. p-A enrichment) protocol and were sequenced on a HiSeq4000 with a 2×75bp data yield and a paired-end mode. The raw data are deposited at the Sequence Read Archive (SRA) and available under the project ID PRJNA909070.

### RNA sequence analysis

RNA-sequencing reads were adapter and quality trimmed using trimmomatic (34) in paired end mode using the following options: ILLUMINACLIP:{adapter.fasta}:2:30:10 LEADING:3 TRAILING:3 SLIDINGWINDOW:4:20 MINLEN:36. Trimmed reads were mapped against the Vienna reference *Nematostella* transcriptome using the Bowtie2 software with default parameters (35,36). Resulting sam files were converted to bam format using the samtools suite (37). Read counts per transcript were estimated by the Salmon software package using default parameters and the -l ISR option (38). Differential analysis of the count data were performed using R and the DESeq2 software package (39,40). For differential gene estimation log-fold change shrinkage was performed before testing difference with a Wald-p-test (betaPrior = TRUE), all genes where the adjusted p-value was lower than α = 0.05 were considered differentially expressed, regardless of fold change.

### Bacteria isolation and culturing

Bacteria were isolated of planula larvae, juveniles and adult polyps. Whole body homogenates were spread out on MB, LB, R2A and count agar plates. Plates were incubated at 4°C, 18°C, 20°C, 30°C or 37°C. Colonies to pick were selected by their morphology in regard to colour, size and shape to exclude redundancy. The goal was to obtain a library of as many bacteria colonizing *Nematostella vectensis* over the whole life cycle as possible under the given culturing conditions. Purified single colonies were transferred into the respective liquid media and saved as either cryostocks or glycerol stocks (10% or 25% final glycerol concentration). If bacteria were regrown for experiments, it was first tried to culture them in MB at 30°C to ensure equal growth conditions. All bacteria used for the mono-association experiments were able to grow on MB.

### Mono-association experiments

Prior to mono-associations, adult polyps were treated with antibiotics in the same matter as for the recolonization experiments. Bacteria for mono-associations were selected for their succession pattern during the recolonization process. Bacteria were grown overnight, diluted in fresh medium and grown to an OD600 of 0.1. Each polyp was recolonized with a calculated OD600 of 0.001 (approx. 50000 cells) of a single bacterial strain in 3mL and incubated at 18°C (n=5). After 24 hours, the medium was exchanged with fresh sterile medium. Seven days after recolonization, polyps were homogenized and spread out on MB plates. Colonies were counted after 3 days of incubation at 18°C.

### Heatmap creation

For the heatmap, ESVs were filtered for their minimal relative abundance during at least one of the four different timepoints of recolonization. The threshold was estimated by calculating the ECDF (empirical cumulative distribution function) and set to 0.6% of the data. Afterwards, the abundance data were normalized to range between 0 and 1 for each sub-heatmap.

### Genome isolation/sequencing/assembly/annotation

Genomic DNA was isolated using the Genomic DNA Purification Kit (Promega) using the protocol for gram positive bacteria. Libraries were prepared using the Nextera DNA Flex Kit (Illumina). Sequencing was performed on an Illumina NextSeq 1500 with a read length of 2*150bp to approximately 60-80X coverage per genome. For assembly, genomic paired-end reads were first trimmed with TrimGalore (41) to remove any remaining adapter sequences and reads shorter than 75 base pairs. Cleaned reads were subsequently assembled into draft genomes with Spades (42) and all-default settings. Finally, for each draft assembly, gene models were annotated using Prokka (43) and its built-in reference database. The raw data are deposited and available under the project ID XXX.

### Metabolic pathway analysis

The inference of bacterial metabolic capacities and the comparison of potential pathway abundances over time was done by metabolic pathway analysis. For the prediction of metabolic pathways gapseq was employed (44). As input served sequence data from newly assembled genomes as well as published genomes from NCBI (**Additional file 2: Table S4**). gapseq was run with default parameters (bitscore threshold of 200) with pathway definitions derived from MetaCyc (45). In addition, other bacteria traits, potentially relevant in host interactions, were inferred using Abricate and the virulence factor database VFDB (46,47). The potential pathway abundances were calculated from genomic capacities and bacterial abundance data. For this means, the relative bacterial abundance for each timepoint, bacterial source, and replicate were computed. Next, for each pathway the sum of relative abundances from all bacteria which were predicted to possess the corresponding pathway were determined. This resulted in relative cumulative pathway abundances that were used to compare changes in metabolic capacities over time.

Pathways associated with early (2d,7d) and late (14d, 28d) time points were summarized to subsystems. Associated pathways were determined by random forest feature selection using Boruta and the importance score of pathways was summed up for each subsystem. Subsystems with an importance score >=0.5 were shown.

## Results

### Adult polyps control initial colonization events

In order to understand the rules underlying the establishment of complex microbial consortia on host tissue, we performed a comparative recolonization experiment on host tissue and inert silicon tubes (**Figure 1A, B**). We used adult polyps of the sea anemone *Nematostella vectensis* that were depleted of their microbiome and sterile silicon tubes to imitate an inactive polyp with an inner (gastrodermic) and an outer (ectodermic) surface. Both, antibiotic-treated polyps und sterile tubes, were recolonized with three different bacterial consortia of larvae (bL), juvenile (bJ) and adult polyps (bA), respectively (**Figure 1A, B**). The three bacterial inocula differed significantly in their composition (**Figure 1C-F. Additional file 1: Figure S1-S2**, pairwise PERMANOVA, pseudo-F value for the polyp experiment: bL-bA 39.94, for bJ-bA 33.99, for bL-bJ 68.52, p and q<0.05; for the tube experiment: bL-bA 90.83, for bJ-bA 43.53, for bL-bJ 104.75 p and q<0.05).

**Figure 1:**
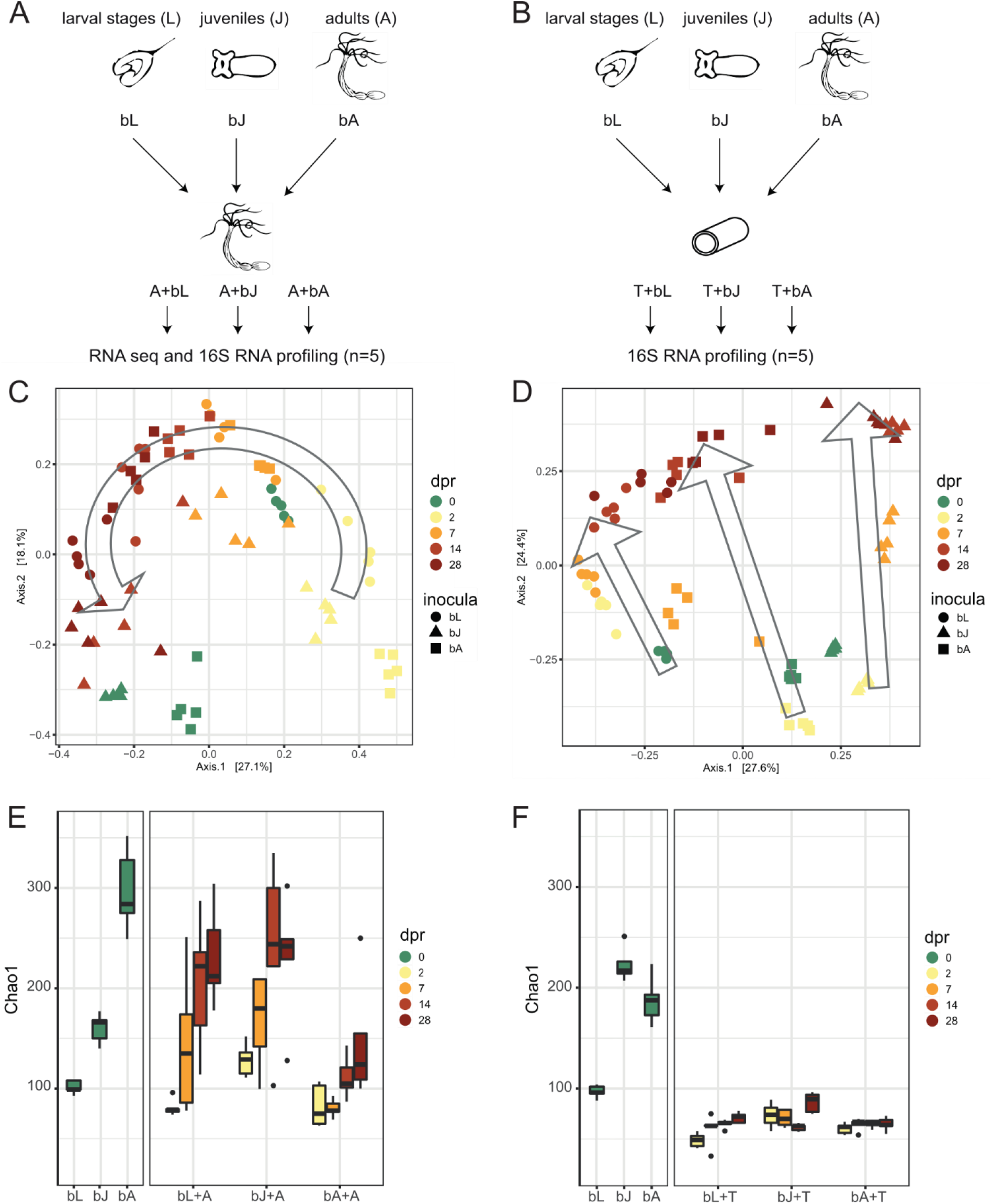
Bacterial recolonization dynamics of germfree polyps and silicone tubes. (A, B) Experimental Setup for recolonization of (A) adult polyps and (B) silicone tubes. Samples were taken from the inocula and 2, 7, 14, and 28 days post recolonization. (C, D) Principal Coordinate Analysis (PCoA) based on the Bray Curtis dissimilarity for the recolonization of (C) polyps and (D) tubes. The arrows indicate the change of the bacterial composition over time. (E, F) Chao1 measure for the recolonization experiment of (E) polyps and (F) tubes. The chao1 meausre of the inocula is shown on the left side, while the change of the chao1 measure over time is shown on the right. The different time points of the recolonization are colour-coded, while the developmental stage from the source of the inoculum is shape-coded. bL=bacteria of Larvae, bJ=bacteria of Juveniles, bA=bacteria of Adults, dpr=days post recolonization.

A comparison of the bacterial community successions on host tissue and silicon tubes revealed significant differences (**Figure 1C, D**). While host recolonization was mainly driven by days post recolonization (dpr) in all three treatments, the recolonization of tubes was influenced by both treatment and time (**Table 1).** Principal coordinate analyses (PCoA) and hierarchical clustering revealed also qualitative differences between host and tube colonization succession (**Figure 1C-D; Additional file 1: Figure S1-S2**). During the colonization of the tubes, initial differences originating from the three different inocula were maintained in the different treatments throughout the experiment (**Figure 1D, Additional file 1: Figure S1**). While principal coordinate 1 (PC1) describes the differences in the bacterial communities of the inocula, PC2 describes the bacterial succession in all three treatments (**Figure 1D**). In contrast, PCoA (**Figure 1C**) and hierarchical clustering (**Additional file 1: Figure S2**) of the bacterial communities recolonizing host tissue revealed a clustering of time points mainly independent of the inocula, even though the beta-diversity distances within time points increased slightly over time (**Additional file 1: Figure S3A**). Already 2 days post recolonization (dpr) the bacterial communities of all three treatments align to each other and show a high similarity to the bacterial communities of larvae (bL) (**Figure 1C**). Interestingly, within the first week of recolonization the similarity to the larvae bacterial (bL) community increases in all three inocula, while 7 dpr the bacterial communities of all three treatments showed the highest similarity to bL (**Figure 1C, Additional file 1: Figure S4A**). Within two weeks of recolonization the bacterial composition of all treatments adjusted to a composition similar to the bacterial composition of juvenile polyps (bJ) (**Figure 1C, Additional file 1: Figure S4B**). 28 dpr the bacterial communities clustered in between the bacterial communities of juvenile and adult polyps. The recolonization pattern of adult polyps, therefore, reflects the pattern of ontogenetic colonization succession (**Figure 1C, Additional file 1: Figure S4C**).

**Table 1.**
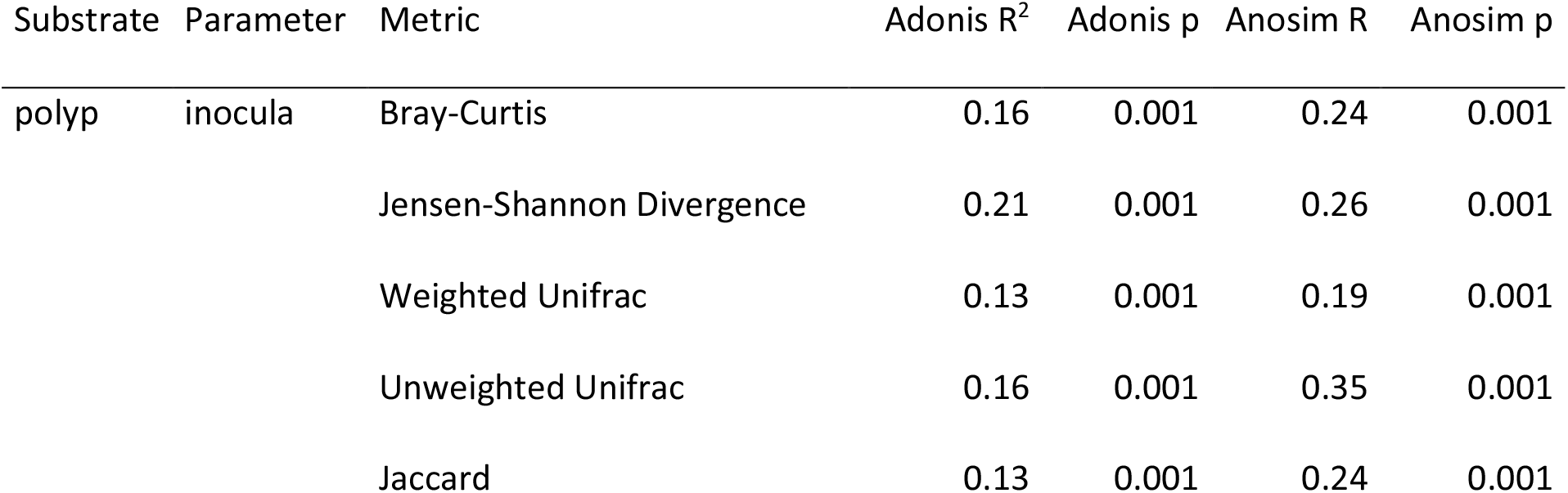
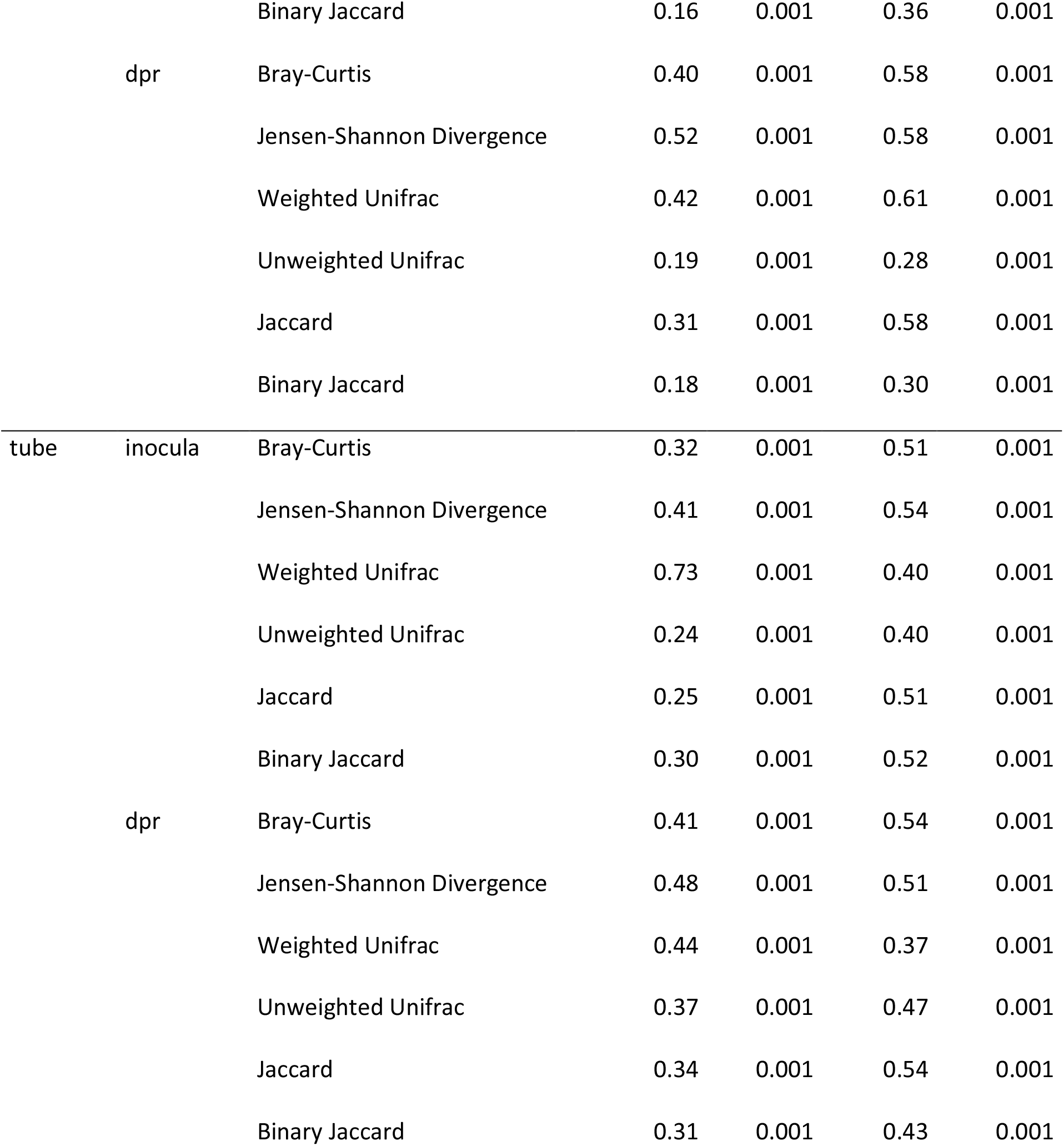
Statistical analysis of the influence of the inocula and the days post recolonization on the recolonization dynamics calculated for six different distance metrices.

| Substrate | Parameter | Metric | Adonis R<sup>2</sup> | Adonis p | Anosim R | Anosim p |
| --- | --- | --- | --- | --- | --- | --- |
| polyp | inocula | Bray-Curtis | 0.16 | 0.001 | 0.24 | 0.001 |
|  |  | Jensen-Shannon Divergence | 0.21 | 0.001 | 0.26 | 0.001 |
|  |  | Weighted Unifrac | 0.13 | 0.001 | 0.19 | 0.001 |
|  |  | Unweighted Unifrac | 0.16 | 0.001 | 0.35 | 0.001 |
|  |  | Jaccard | 0.13 | 0.001 | 0.24 | 0.001 |
|  |  | Binary Jaccard | 0.16 | 0.001 | 0.36 | 0.001 |
|  | dpr | Bray-Curtis | 0.40 | 0.001 | 0.58 | 0.001 |
|  |  | Jensen-Shannon Divergence | 0.52 | 0.001 | 0.58 | 0.001 |
|  |  | Weighted Unifrac | 0.42 | 0.001 | 0.61 | 0.001 |
|  |  | Unweighted Unifrac | 0.19 | 0.001 | 0.28 | 0.001 |
|  |  | Jaccard | 0.31 | 0.001 | 0.58 | 0.001 |
|  |  | Binary Jaccard | 0.18 | 0.001 | 0.30 | 0.001 |
| tube | inocula | Bray-Curtis | 0.32 | 0.001 | 0.51 | 0.001 |
|  |  | Jensen-Shannon Divergence | 0.41 | 0.001 | 0.54 | 0.001 |
|  |  | Weighted Unifrac | 0.73 | 0.001 | 0.40 | 0.001 |
|  |  | Unweighted Unifrac | 0.24 | 0.001 | 0.40 | 0.001 |
|  |  | Jaccard | 0.25 | 0.001 | 0.51 | 0.001 |
|  |  | Binary Jaccard | 0.30 | 0.001 | 0.52 | 0.001 |
|  | dpr | Bray-Curtis | 0.41 | 0.001 | 0.54 | 0.001 |
|  |  | Jensen-Shannon Divergence | 0.48 | 0.001 | 0.51 | 0.001 |
|  |  | Weighted Unifrac | 0.44 | 0.001 | 0.37 | 0.001 |
|  |  | Unweighted Unifrac | 0.37 | 0.001 | 0.47 | 0.001 |
|  |  | Jaccard | 0.34 | 0.001 | 0.54 | 0.001 |
|  |  | Binary Jaccard | 0.31 | 0.001 | 0.43 | 0.001 |

Analysis of the degree of restructuring of bacterial communities in the different treatments showed that the bacterial communities of polyps recolonized with bacteria of adult polyps were the most restructured (**Additional file 1: Figure S3B**). In contrast, the bacterial community of animals recolonized with bacteria from larvae exhibited the lowest degree of restructuring (Anosim R=0.2364, p<0.001; **Additional file 1: Figure S3B**). However, within 28 dpr the three treatments did not approach the identity of the adult bacterial inoculum completely (**Figure 1C, Additional file 1: Figure S4C**). Compared with the wild-type control polyps, which spent the same time in sterile medium without food as the treatment polyps, the treatment polyps approached the wild-type controls after 28 dpr (**Additional file 1: Figure S5**). This suggests that the difference between the adult microbiota and those recolonized for 28 dpr (**Figure 1C**) may be due to starvation.

The comparisons of the alpha-diversity also revealed significant qualitative differences. While the bacterial diversity increased during the bacterial succession on the host and approach the level of the adult bacterial inoculum after four weeks in all treatments (**Figure 1E**), the bacterial diversity on the tubes remained stable on a low level (**Figure 1F**). The changes in absolute bacterial abundance in the two experiments were also opposite. While bacterial abundance on the host tissue decreased over the course of the experiment (**Additional file 1: Figure S6A**), bacterial abundance on the tube increased within the four-week experimental period (**Additional file 1: Figure S6B**).

These results suggest that the mechanisms controlling bacterial colonization of host tissue and inert silicone tubes differ significantly. Whereas on the silicone tubes the inocula determined the initial colonization events and thus the subsequent colonization, the initial colonization events on the host tissue were mainly independent of the inocula. Here, early colonization events in all treatments were characterized by similar bacterial communities corresponding to the microbiota of early life stages and subsequent colonization resembled ontogenetic colonization pattern. Thus, we conclude that the initial colonization events appear to be strongly influenced by the host but not by the tube, while the subsequent bacteria-bacteria interactions are the main cause of the observed bacterial succession in both host tissue and the tube.

### Recolonization successions resemble ontogenetic colonization sequence

To determine whether the similarity in polyp recolonization between the three different treatments was due to similar bacterial groups or the same initial colonizers, and to identify bacteria that might act as drivers of the observed bacterial succession, we examined bacterial succession at the exact sequence variant (ESV) level. Therefore, we compared the abundances of the ESVs that are present in at least one of the three inocula with their abundances over the recolonization process (**Figure 2A, Additional file 2: Table S1**). For this, we sorted for the ESVs with a minimum relative abundance of 0.6% during at least one of the four different timepoints of recolonization (61 ESVs). In the first column of the heat map the relative abundance of the selected ESV in the three inocula, bL, bJ and bA, are indicated, illustrating three distinct groups of ESVs characterizing the three different inocula. The three subsequent columns illustrate the relative abundance of these ESVs in the three different recolonization experiments, adult polyps (A) recolonized with bacteria of larvae (+bL), of juvenile polyps (+bJ) and adult polyps (+bA) (**Figure 2**).

**Figure 2:**
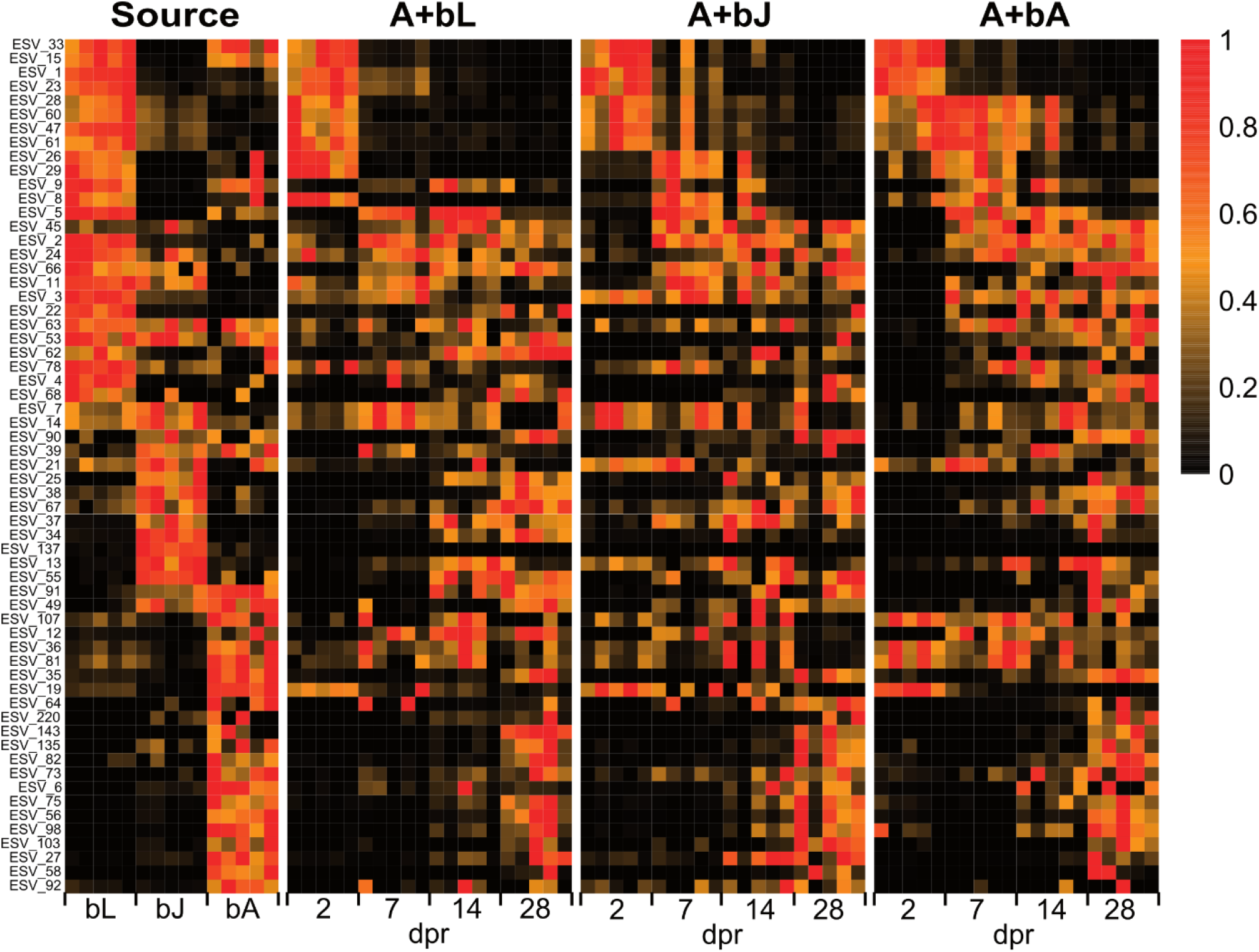
Recolonization dynamics on single ESV level for ESVs with a minimal relative abundance of 0.6% during at least one of the four different timepoints of recolonization (61 ESVs). The left column represents ESVs present in the microbiome of larvae (bL), juveniles (bJ) and adults (bA), while the next three columns represent the temporal appearance of the 61 ESVs during the recolonization with bacteria isolated from larvae (A+bL), from juveniles (A+bJ), and from adults (A+bA). The five replicates per treatment are shown as separate cells within the columns.

Interestingly, all three treatments show a similar pattern as seen in the inocula themselves. 2 and 7 dpr the most common ESVs are larval specific in all three treatment (**Figure 2**). After 14 dpr, the most common ESVs are specific to larval and juvenile stages, while the relative abundance of some larval-specific ESVs is already decreasing. After 28 dpr, adult-specific ESVs emerge in the community of all three treatments (**Figure 2**). Therefore, the most abundant ESVs during the early recolonization process are specific for larval developmental stages, while the most abundant ESVs during the late recolonization are specific for late developmental stages. We conclude from this that the first colonization events are initiated by larval-specific bacteria and that successively the community composition approaches the identity of the bacterial community of adult polyps. In addition, we infer that these mechanisms are deterministic instead of stochastic for its consistency through all three treatments and replicates.

### Early colonizers show a higher capability of colonization compared to late colonizers

We performed a culturing approach to test the hypothesis that early colonizing bacteria can readily colonize *Nematostella*, while late-emerging bacteria only colonize poorly in mono-association. We isolated 161 bacteria from different developmental stages of *Nematostella* by plating tissue homogenates on three different bacterial media (**Additional file 2: Table S2**). These isolates belong to a range of Alpha-, Beta- and Gammaproteobacteria, as well as Actinobacteria, Firmicutes and one Bacteroidetes strains (**Additional file 2: Table S2**). However, we were unsuccessful in culturing Deltaproteobacteria, Planctomycetes and Spirochaetes.

These isolates were mapped to the ESVs from the recolonization experiment and based on the relative abundance of the ESVs during the recolonization process, they were classified into “early” and “late” colonizers (**Figure 2**, **Additional file 2: Table S2**). If the abundance of an ESV in the inocula did not match the abundance during the recolonization based on the heatmap (**Figure 2**), the ESV was assigned based on recolonization pattern.

To test the hypothesis that early colonizers have a higher ability to recolonize adult polyps compared to late colonizers, we performed mono-association experiments. For this, we selected five bacterial isolates representing ESVs that recolonize early during the recolonization experiment, and five bacterial isolates representing ESVs that recolonize late during the recolonization experiment.

While all ten bacterial strains were able to colonize on germfree polyps, early bacteria colonized *Nematostella* with a significantly higher density than late-appearing bacteria (Kruskal-Wallis chi-squared=16.528, p<0.0001, **Figure 3**). Thus, initial colonization appears to be controlled by the promotion or inhibition of specific bacterial strains, which may be driven by metabolic dependencies or host-controlled mechanisms.

**Figure 3:**
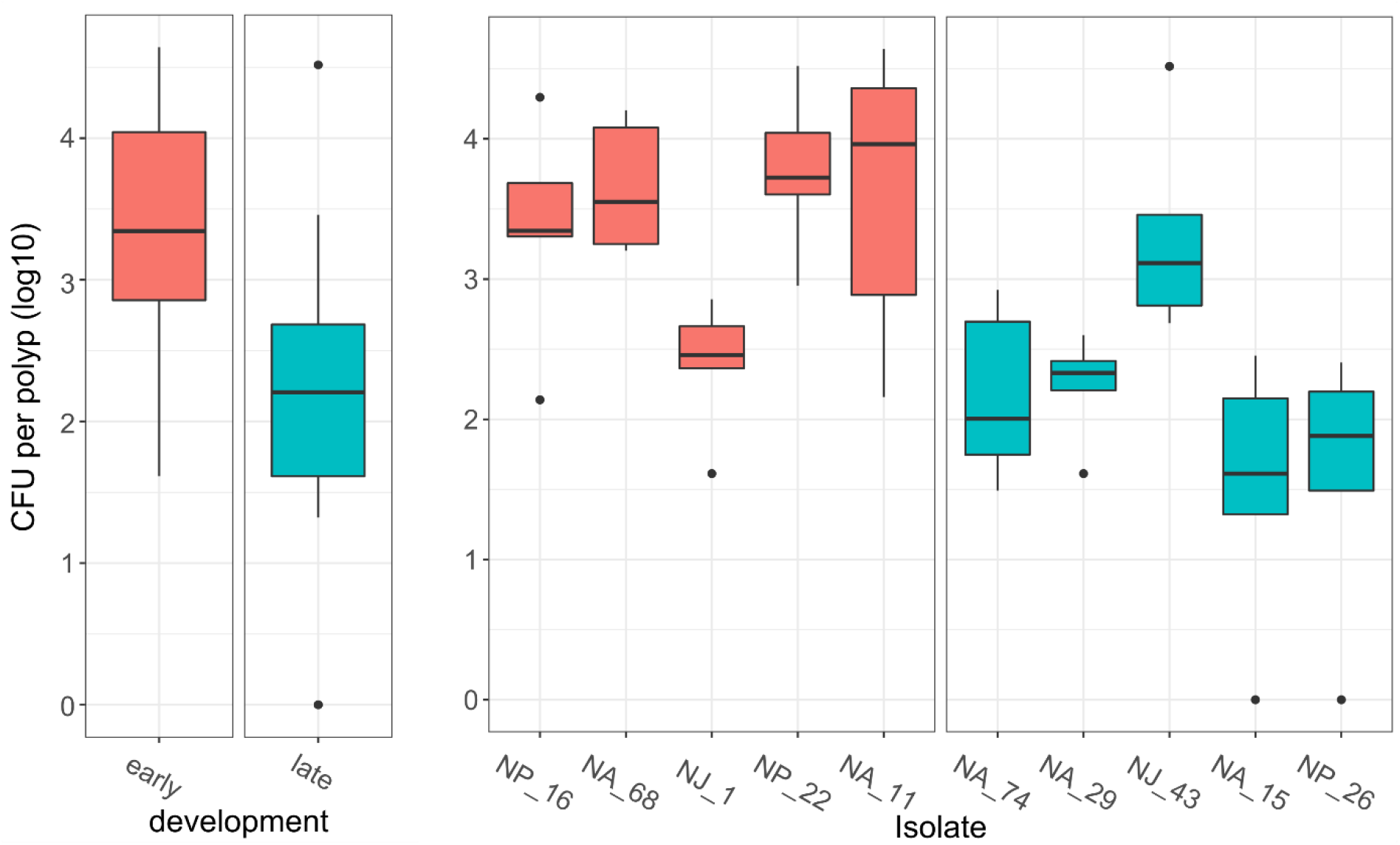
Mono-associations of germfree adult polyps with single bacterial strains. Counted CFU per polyp after recolonization with single bacteria. Isolates were classified as early- or late-appearing depending on their appearance during early or late recolonization. Polyps were recolonized with single bacterial isolates for seven days before polyps were homogenized and spread on MB plates. Colonies were counted after three days of incubation (n=5). On the left were all early- or all late-appearing bacteria pooled. On the right bacteria are shown separately. Data were log10-transformed.

### Metabolic capabilities reflect recolonization pattern

We reconstructed the metabolic networks of bacterial colonizers to estimate the metabolic potential relevant to colonization and species interactions. The metabolic networks of 31 sequenced isolates (**Additional file 2: Table S3**) and additional 125 publicly available genomes (**Additional file 2: Table S4**) were obtained, whereby the selected 16S rRNA genes of the publicly available genomes matched with ESVs from the colonization process by at least 97%. Metabolic networks contain the predicted enzymatic reactions and pathways of an organism and were used to compare metabolic capabilities. We combined the 16S rRNA abundance data with predicted metabolic networks to derive potential pathway abundances for each time point during recolonization.

First, we investigated if the unique bacterial colonization succession can be found again on the metabolic level. **Figure 4A** shows the variance of metabolic pathway abundances between samples during colonization as a PCA plot. We found that pathway abundances indeed reflected the observed recolonization pattern, indicated by a separation in dimension one of the PCA across time points of colonization. Pathways that contributed most to dimension one were pathways involved in biosynthesis (e.g. cofactor, amino acids, lipids), degradation (e.g. carbohydrates), and energy metabolism (**Additional file 2: Table S5**). Given the reappearing pattern on the metabolic level, we performed feature extraction by random forests to find pathways associated with early (days=2,7) and late (days=14,28) colonizers. We identified 57 pathways reported consistently in repeated feature extractions, and these pathways correctly classified all samples into early or late stages with an accuracy of 93% in k-fold cross-validation.

**Figure 4:**
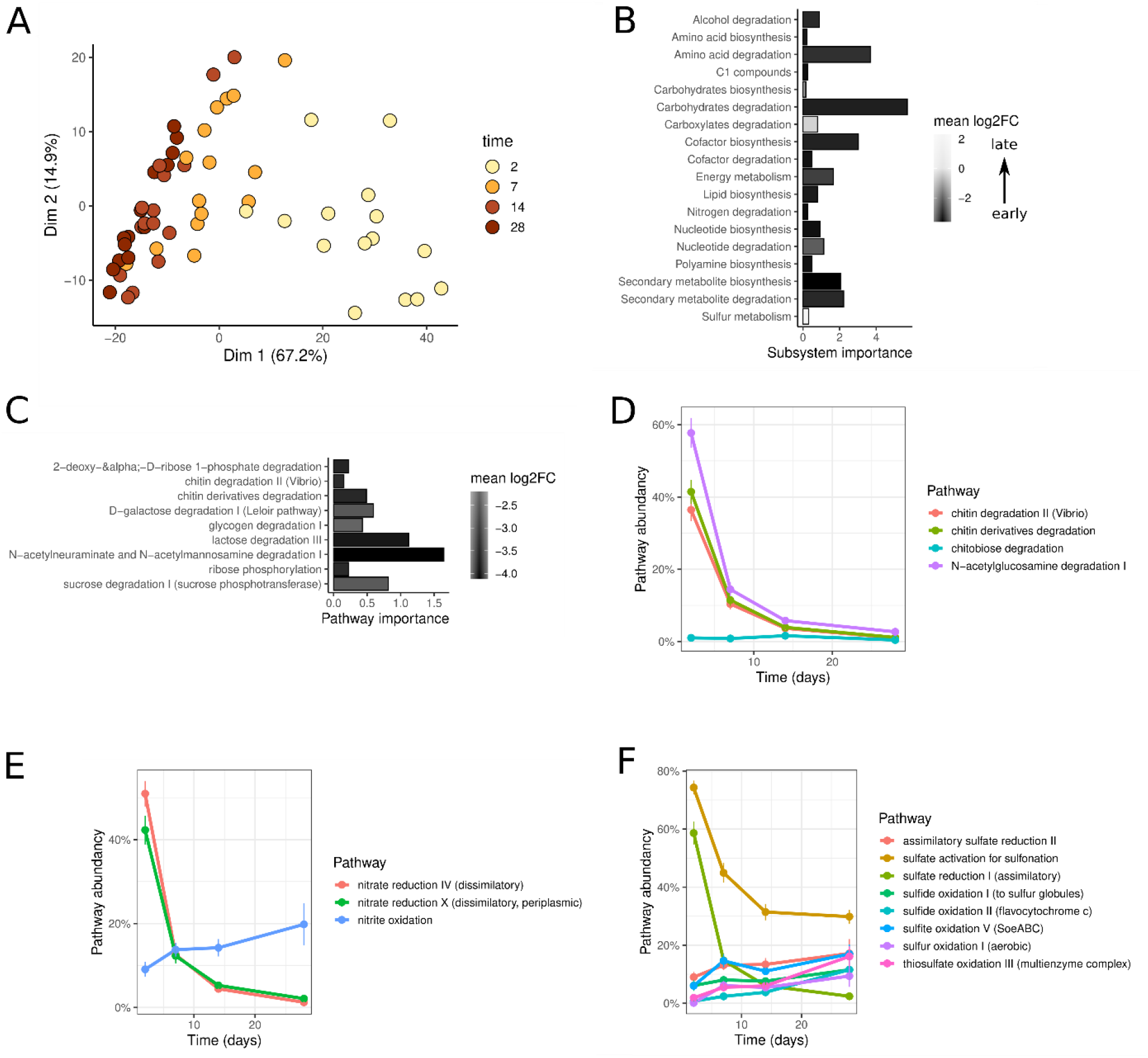
Reconstructions of metabolic networks during bacterial succession A) Principal component analysis (PCA) of the metabolic capabilities of the recolonization samples. Each sample contains the metabolic pathway abundances that were derived from inferred metabolic pathways combined with 16S rRNA abundance data. Colors indicate the time point of the sample. B) Pathways associated with early (2d,7d) and late (14d, 28d) time points were summarized to subsystems. The filling indicates the early vs. late colonizers mean log2 fold change of the pathway abundances at early and late time points. C) Carbohydrate degradation pathways separating early vs. late colonizers from random forest feature selection. The log2 fold change of mean pathway abundances at early (2d,7d) and late (14d, 28d) time points is shown together with the importance score from random forest feature selection (Boruta). D) Time series of chitin degradation associated pathway abundances. The pathway abundance indicates the distribution of pathways among colonizing bacteria (based on 16S relative abundances). Time series of pathway abundances for E) nitrogen and F) sulfur cycle associated pathways.

When pathways were summarized into subsystems, the importance of carbohydrate and amino acid degradation was highest (**Figure 4B**). Again carbohydrate degradation showed the most remarkable changes with a mean log2 fold change higher than −3. Among the identified carbohydrate degradation pathways, polysaccharide degradation (chitin, glycogen), and sugar catabolism (ribose, galactose, lactose, ribose, sucrose) were dominant (**Figure 4C**).

Interestingly, we found degradation of chitin and its derivatives among the list of carbohydrate degradation pathways. The pathway abundances of chitin degradation related pathways showed high variance over time, reaching the maximum of 40-60% in early time points (**Figure 4D**). The degradation of chitin into monomers of N-acetyl-glucosamine could be accompanied by further utilization. In line with this, the degradation of N-acetyl-glucosamine showed higher abundances also at later time points. (**Figure 4D**).

In addition, potential bacteria-bacteria interactions involved in bacterial succession were identified by the investigation of complementary pathways and by comparing the changes in their abundances. Nitrate reduction was among the pathways identified by random forest feature selection (subsystem nitrogen degradation in **Figure 4B**, **Additional file 1: Figure S7**). Nitrogen cycling pathways showed a potential link of early nitrate to nitrite reduction with later nitrite oxidation (**Figure 4E**). Similarly, feature selection identified hydrogen sulfide oxidation for later colonizers (subsystem sulfur metabolism in **Figure 4B**). Moreover, sulfur cycling pathways suggested the early reduction of sulfate followed by later oxidation (**Figure 4F**).

### *Nematostella* shows a common transcriptomic response to bacterial recolonization including chitin synthesis

To identify host mechanisms and functions that might be involved in the selection of early colonizers, we analyzed the common host response to bacterial recolonization. Therefore, we extracted and sequenced the host’s mRNA 2 dpr and compared the response to the three different inocula to each other and to the germfree controls (**Figure 5A**). In total, 4103 genes were differentially regulated in recolonized animals in comparison to germfree animals, which represent almost 16% of the whole transcriptome (25729 genes) (**Figure 5B**). Analyzing the host responses to the three inocula, it is notable that animals responded most strongly to the adult inoculum. In total, 426 genes were differentially regulated in response to the adult inoculum, in contrast to 82 and 91 genes in response to the juvenile and larval inoculum, respectively. This result agrees well with the observation that the microbiota of adult polyps undergoes the greatest restructuring during recolonization (**Figure 1**) and that most likely the host is controlling these early colonization events.

**Figure 5:**
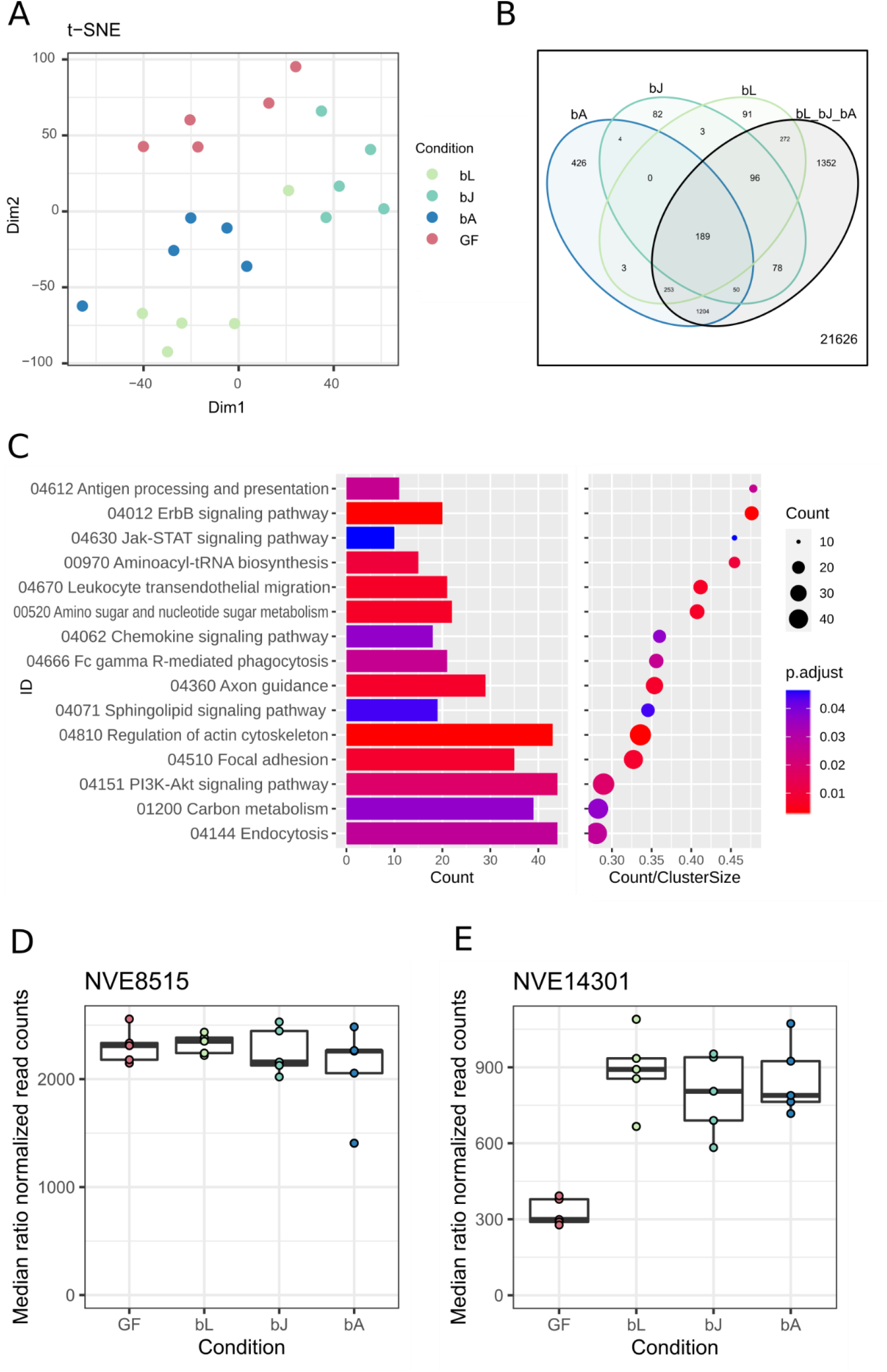
Transcriptomic analysis of recolonized adult polyps 2 days post recolonization. (A) t-SNE plot of the sequenced samples clustering according to their treatment. GF=germfree. (B) Venn diagram of regulated genes in all three treatments and their overlaps. To increase the statistical power, the three comparisons bL vs GF, bJ vs GF and bA vs GF were separately done to the fourth comparison of bL-bJ-bA vs GF. This way, 1352 more genes could be found that are differentially regulated in all three treatments. (C) Regulation of KEGG clusters of all three treatments versus germfree polyps. The barplots show the counts of the genes belonging into the different KEGG clusters, while the dots represent the ratio of the counts to the size of the cluster. (D) Normalized read counts of the two chitin synthase genes NVE8515 and NVE14301 in Nematostella, 2 days post recolonization. (A) Normalized read counts for NVE8515 in recolonized animals compared to germfree (GF) animals. (B) Normalized read counts for NVE14301 in recolonized animals compared to germfreec animals (log2fc=2.04, p<0.001).

189 genes are upregulated in all three treatments and therefore represent genes that are generally upregulated upon contact and eventually colonization by commensal bacteria. They belong to a variety of KEGG categories (**Figure 5C, Additional file 2: Table S6**). ErbB signaling and Jak-STAT signaling transduce signals through the PI3K-Akt pathway to influence cell proliferation, differentiation, motility and survival. Fc gamma R-mediated phagocytosis, regulation of actin skeleton and endocytosis are all involved in engulfment of particles of various sizes. Antigen processing and presentation, leukocyte transendothelial migration and focal adhesion could indicate a dynamic immune cell response. The enrichment of the KEGG clusters of amino sugar and nucleotide sugar metabolism and carbon metabolism pinpoints towards a mechanism involved in carbohydrate metabolism. Interestingly, among the commonly regulated genes we found within the Top 20 most upregulated genes one of the two chitin synthase genes present in *Nematostella* (**Figure 5D, Additional file 2: Table S6**), while a second chitin synthase was not differentially expressed (**Figure 5E,** NVE8515). The gene NVE14301 is upregulated consistently in response to all bacterial recolonization treatments (**Figure 5D**). The coincidence of the high prevalence of chitin degraders among early colonizers and the upregulation of the host’s carbon metabolism, specifically a chitin synthase, suggests that chitin might be essential for the interaction between host and early colonizers.

## Discussion

### Early colonization events are determined by the host and not by priority effects

Analysing the host transcriptomic response to the three different bacterial consortia revealed that *Nematostella* reacts strongly to the bacterial recolonization. Remarkably, the strongest response by far was exhibited by the adult polyps that were recolonized with adult bacteria. Looking at the 16S phylogenetic analysis, the community composition resets to a larval-like community, so the adult inoculum must undergo the greatest restructuring. The highest number of regulated genes in these animals suggest that this “reset” is not a bacterial driven process but mainly a host driven one. This argument is supported by the recolonization of inert silicone tubes. Here, the recolonization succession was mostly dependent on the inoculum and the recolonizations with the three inocula remained separated.

The common response of the polyps to all three inocula suggests how *Nematostella* generally responds to, interacts with and selects bacterial colonizers. As several pathways regarding phagocytosis are upregulated in combination with signalling pathways involved in cell proliferation and immune cell migration, this suggests that the early transcriptomic response to the recolonization process is a response of the cellular innate immune system. Through its innate immune system, the host can influence and regulate the bacterial communities in a diverse manner. It does not only serve as a defence barrier against pathogens, but also regulates the composition of commensal microbes via e.g. MyD88-dependent pathways (48,49) or by the production of AMPs (50,51). In *Nematostella*, there’s growing evidence that nematosomes, small motile multicellular bodies in the gastric cavity, are part of the cellular innate immune system in *Nematostella* (52,53). They co-express components of the TLR signalling pathway, as TLR and NF-kB (53), and are able to phagocytose foreign particles and bacteria (52,54). As the transcriptomic response to recolonization is dominated by cell proliferation, phagocytosis and motile immune cell migration, we here further support the hypothesis of nematosomes as part of the innate immune system. Future studies will reveal whether nematosomes, in the form of free-floating immune cell structures, have the ability to selectively phagocytose bacteria and thereby influence colonization of the adult polyp. Differential phagocytosis is already known in the squid-vibrio system, where haemocytes are able to differentiate between the squid’s preferred bacterial symbiont *Vibrio fisheri* and other bacteria of the *Vibrio* genus (55). In this study they show that phagocytosis of *V. fisheri* was reduced by pre-exposure of haemocytes to the bacteria, and by the presence of the outer membrane protein OmpU on *V. fisheri*. In a leech model, it is shown that the disruption of the type III secretion system in *Aeromonas veronii* made them vulnerable for phagocytosis by the leech’s macrophage-like cells, while also reducing its pathogenicity in a mouse septicemia model (56).

In regard to the elevated microbial metabolic potential to degrade chitin during the first two days, one of the most interesting upregulated host genes is a chitin synthase. Although it has already been known for several years that *Nematostella* possesses at least two genes for chitin synthesis (57), there is just emerging evidence that soft-bodied anemones also express chitin synthase genes. The expression strength indicates a mechanism where chitin is continuously produced while bacteria are present but its production halts if bacteria are missing (**Figure 5E**).

In parallel to the increased expression of a chitin-producing enzyme, early colonizing bacteria had an increased capacity to degrade chitin. Pathways from carbohydrate degradation were most distinguished between early and late colonizers and among them chitin pathways were prominent. The host’s production of chitin, therefore, seemed to be accompanied by microbial utilization.

In general, chitin is widely available in the ocean and chitin degradation activity has been detected for many marine bacteria (58,59). In addition, micro-particles of chitin have been shown to enable the community assembly of free-living seawater bacteria (60). In the context of host-microbiome associations, host-produced chitin is known to modulate immune response (61). It has been proposed to enable gut compartmentalization and thus permit barrier immunity from which the mucus layer and its microbial colonization might have been evolved (62). From this we concluded that chitin might also play a central role in host-microbiota interactions in *Nematostella* and potentially also in the succession. For cnidarians, the functionality of chitin synthases has been described (57). We hypothesize that host-produced chitin creates a distinct niche that allows chitin-degrading bacteria to flourish and causes the observed succession dynamics.

Commonly, the processes influencing the community assembly can be a combination of deterministic and stochastic processes (9). Deterministic processes include mechanisms such as the host’s genetic background, its immune system, nutrition, metabolic prerequisites, or environmental factors. Highly deterministic effects are observed in systems such as the *Vibrio* squid system, in which the squid selects *V. fisheri* that is induced to colonize by the production of chemoattractants such as chitobiose and nitric oxide and by attraction via motile cilia (63–65). Here, the host has complete control over bacterial colonizers. Stochastic processes include priority effects or passive dispersal. In systems where stochasticity is more critical, e.g., due to priority effects, perturbations in microbial composition are observed long during ontogeny, if not into adulthood. Consequently, children born via C-section exhibit a different microbiome than those born vaginally (66), and high levels of hospital pathogens can colonize the infant’s gut, disrupting the transmission of Bacteroides/Bifidobacterium and other commensals. (67).

In contrast, our data indicate that priority effects do not play a significant role in *Nematostella*, as the recolonization dynamics are mainly independent of the inoculum. Similarly, Mortzfeld et al. 2015 stated that the developmental age of the host is the main driving force of the *Nematostella* microbiome (14). However, here we could show that not host ontogeny, but host niches and interactions are driving the community composition as we performed experiments on animals which already completed their development.

### Bacterial succession depends mainly on bacteria-bacteria interactions rather than host development

Once the host has shaped the initial microbial community, microbial forces show a stronger influence on community succession. We observed consistent dynamics up to the establishment of the adult microbiota independent of host development. Recolonization of adult polyps resulted in a microbial community resembling the community typical of the larval stage, followed by shifts toward a juvenile and adult microbiota. Microbial taxa found at later time points therefore followed a non-random trend. Our mono-association experiments showed higher recolonization success of early-colonizing bacteria compared to late-colonizing bacteria, indicating a mechanism promoting a faster settlement of early colonizers.

Because the early colonizing bacteria are not necessarily the bacteria found in large numbers in the adult polyps, they may fit the definition of a keystone species that is present in small numbers but plays a critical role in maintaining community organization and diversity (68). Especially in ecological systems, keystone species and foundation species are essential for subsequent colonization e.g. in seaweed forests or in habitats after disturbance (69,70). This can imply a niche differentiation or cross feeding events. Datta et al 2016 show that when chitin-covered magnetic beads are submerged in natural marine seawater, the colonization of these beads is mostly determined by the metabolic potential of the bacteria and can be divided into three parts (60). The first bacteria to settle are specialized in attachment; the second are specialized in metabolizing chitin. The third and last wave of bacteria are specialized in feeding on secondary metabolites of chitin degradation. Similarly, we found chitin followed by chitin derivatives degradation for the *Nematostella* microbiome, potentially supporting similar bacteria-bacteria interactions. Interestingly, the alpha-diversity over time of these colonized beads showed a similar pattern as the alpha-diversity in naturally developing *Nematostella* polyps (14). The authors hypothesized that this strong drop of alpha diversity shortly after hatching is an effect of metamorphosis and may represent a bottleneck during development. However, the data of Datta et al. and the results presented here are more suggestive of a metabolic bottleneck in the microbiome itself.

Another ambiguity lies in the coherence of the juvenile and the adult microbiome. It is debatable if the juvenile state is an intermediate state before the mature (adult) state is reached, or if it is an alternative state of the microbiome. The insecurity about that arises firstly from the results of the recolonization, where after one month of recolonization, the adult polyps still did not reach the state of the adult inoculum (**Figure 1C**), and secondly from the data of wildtype animals that were starved throughout the experiment, whose microbiome converged to the juvenile microbiome over time (**Additional file 1: Figure S5**). It may be that starvation has a rejuvenating effect on the microbiome. It was shown in several species like fruit flies, rats and nematodes that caloric or dietary restriction extends the life span by probably downregulating insulin und insulin-like signaling, the amino signaling target of rapamycin (TOR)-S6 kinase pathway, and the glucose signaling Ras-protein kinase A (PKA) pathway (71–73). The microbiome can also pose a positive influence on longevity by integrating cues from diet which have been shown with a drug-nutrient-microbiome screen (74). Therefore, we hypothesize that the nutritional state of the polyp influences the microbial interactions towards a rejuvenated microbiome.

We identified further potential bacteria-bacteria interactions influencing the observed dynamic when investigating metabolic cycles. Early colonizers showed an increased capacity to reduce nitrate and sulfate, whereas later colonizing species could oxidize the reduced compounds (nitrite, sulfite, H2S). However, as nitrate and sulfate reduction are mostly carried out by anaerobic bacteria, there’s an indication that *Nematostella* provides anaerobic niches. In stony corals, extreme diel fluctuations of oxygen in the vicinity of the polyps, as well as anaerobic nitrate reduction could be shown (75,76). Oxidation of reduced nitrogen compounds is also a very common process found in coral reefs (77). Therefore, we propose that interacting reduction-oxidation pathways are important drivers of the bacterial succession dynamics.

## Conclusion

In summary, we uncovered a distinct colonization pattern for the microbiota of *Nematostella* that consistently resulted in very similar bacterial succession in recolonization experiments, regardless of the initial community (**Figure 6**).

**Figure 6:**
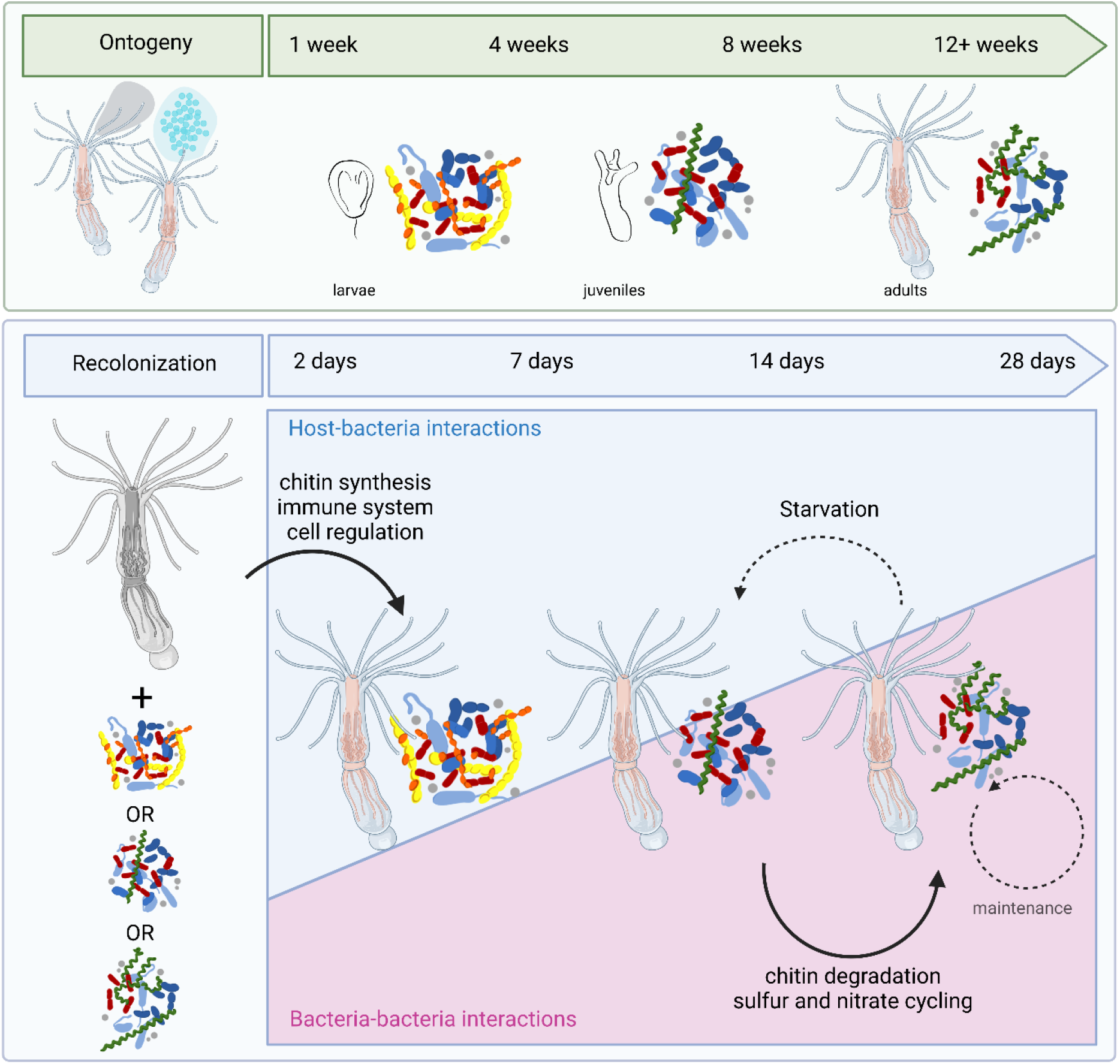
Comparison of the bacterial succession on Nematostella during ontogeny and during recolonization. During ontogeny, larvae, juvenile polyps and adult polyps possess distinct bacterial communities. During recolonization of adult polyps, this bacterial colonization pattern occurring during natural development is recapitulated, independent of the developmental stage from which the bacterial inoculum was isolated. While the bacterial successions during ontogeny take around 3 months, the bacterial successions during recolonization take roughly four weeks. While initial selection of bacterial colonizers during recolonization is mainly directed by the host, subsequent bacterial succession and maintenance are mainly controlled by bacteria-bacteria interactions. Starvation of the host results in a rejuvenation of the microbiome towards a juvenile state. Image created in Biorender. This colonization pattern recapitulates the colonization pattern occurring during ontogeny, however, in a shorter time frame. As the bacterial successions are independent of the initial community, we conclude that in a marine model system, the establishment of colonization is shaped by the host and not by priority effects. Subsequent bacterial succession is mainly determined by bacteria-bacteria interactions, which show a subsequent chitin degradation as well as sulfur and nitrate cycling pathway enrichment during recolonization.

## Availability of data and materials

“The datasets supporting the conclusions of this article are available in the Sequence Read Archive (SRA) under the accession numbers PRJNA902551 (https://www.ncbi.nlm.nih.gov/bioproject/PRJNA902551) and PRJNA909070 (https://www.ncbi.nlm.nih.gov/bioproject/909070).

## Additional material

**Additional file 1:** Supplementary figures, SuppFigures.docx.

**Figure S1:** Analysis of the bacterial recolonization dynamics based on 16S upon recolonization of silicone tubes over the course of one month. **Figure S2:** Analysis of the bacterial recolonization dynamics based on 16S upon recolonization of adult polyps over the course of one month. **Figure S3:** Bray-Curtis Dissimilarity Ranks of the bacterial community depending on the time and on the inocula. **Figure S4:** Bray Curtis distance of the bacterial communities on recolonized animals over time in comparison to the inocula. **Figure S5:** Recolonization dynamics of germfree polyps over the course of one month. **Figure S6:** Absolute bacterial load of the polyps (A) and silicone tubes (B) over the course of the recolonization process.

**Additional file 1**: Supplementary tables, SuppTable.xlsx.

**Table S1**: ESVs with shortened ESV number and 97% cluster to which they belong. **Table S2**: Bacterial strains isolated from Nematostella vectensis with phylogeny according to GenBank Accession number, and ESV names**. Table S3**: Bacterial strains from which genomes were sequenced, with developmental stage from which they were isolated, phylogeny and ncbi classification. **Table S4**: Clusters with the genomes (self-sequenced or downloaded from the ncbi database) which were used for the metabolic potential analysis. **Table S5**: Pathways contributing the most to the separation on dimension 1 in the PCA showing the metabolic capabilities during recolonization (Figure 4A). **Table S6**: Top20 upregulated genes upon recolonization, independent of the inoculum. The upregulation is shown as the Log2 fold change (Log2FC). The p-value is smaller than 5.204e-11 for all of these 20 genes and therefore not separately stated.

## Acknowledgements

We thank Katja Cloppenborg-Schmidt for preparing the 16S rRNA gene library. We thank Peter Deines for the constructive discussions on the tube experiments.

## Funding

This work was supported by DFG CRC grant 1182 “Origin and Function of Metaorganisms” (Project B1, A1, Z2; Z3 and INF). NGS was carried out at the Competence Centre for Genomic Analysis (Kiel) within the CRC 1182 project Z3.

## Author information

Authors and affiliations

**Institute for Zoology and Organismic Interactions, HHU Düsseldorf, 40225 Düsseldorf, Germany**

Hanna Domin, Sebastian Fraune, Gabriela Maria Fuentes Reyes, Lucy Saueressig

**Research Group Medical Systems Biology, Institute of Experimental Medicine, CAU Kiel, 24105 Kiel, Germany**

Johannes Zimmermann, Jan Taubenheim, Christoph Kaleta

**Institute for General Microbiology, CAU Kiel, 24105 Kiel, Germany**

Daniela Prasse, Ruth Anne Schmitz

**Sysmex Inostics, 20251 Hamburg, Germany**

Daniela Prasse

**Institute for Clinical Molecular Biology, CAU Kiel, 24105 Kiel, Germany**

Marc Höppner

**RD3 Marine Symbioses, GEOMAR Helmholtz Centre for Ocean Research, 24105 Kiel, Germany**

Ute Hentschel

**CAU Kiel, 24105 Kiel, Germany**

Ute Hentschel

## Contributions

HD, JZ, CK, UH and SF conceived and designed the study. HD, LS and GFR conducted the experiments. DP and RAS isolated and provided the bacterial library. HD, JZ, JT and MH analysed the data. HD, SF and JZ wrote the manuscript. All authors revised the manuscript and read and approved the final manuscript.

## Corresponding author

Correspondence to Sebastian Fraune

## Ethics declaration

Ethics approval and consent to participate

Not applicable.

## Consent for publication

Not applicable.

## Competing interests

The authors declare that they have no competing interests.

## Notes

### Competing Interest Statement

The authors have declared no competing interest.

