## Supplementary Figures for "Bacterial colonizers of *Nematostella vectensis* are initially selected by the host before interactions between bacteria determine further succession"

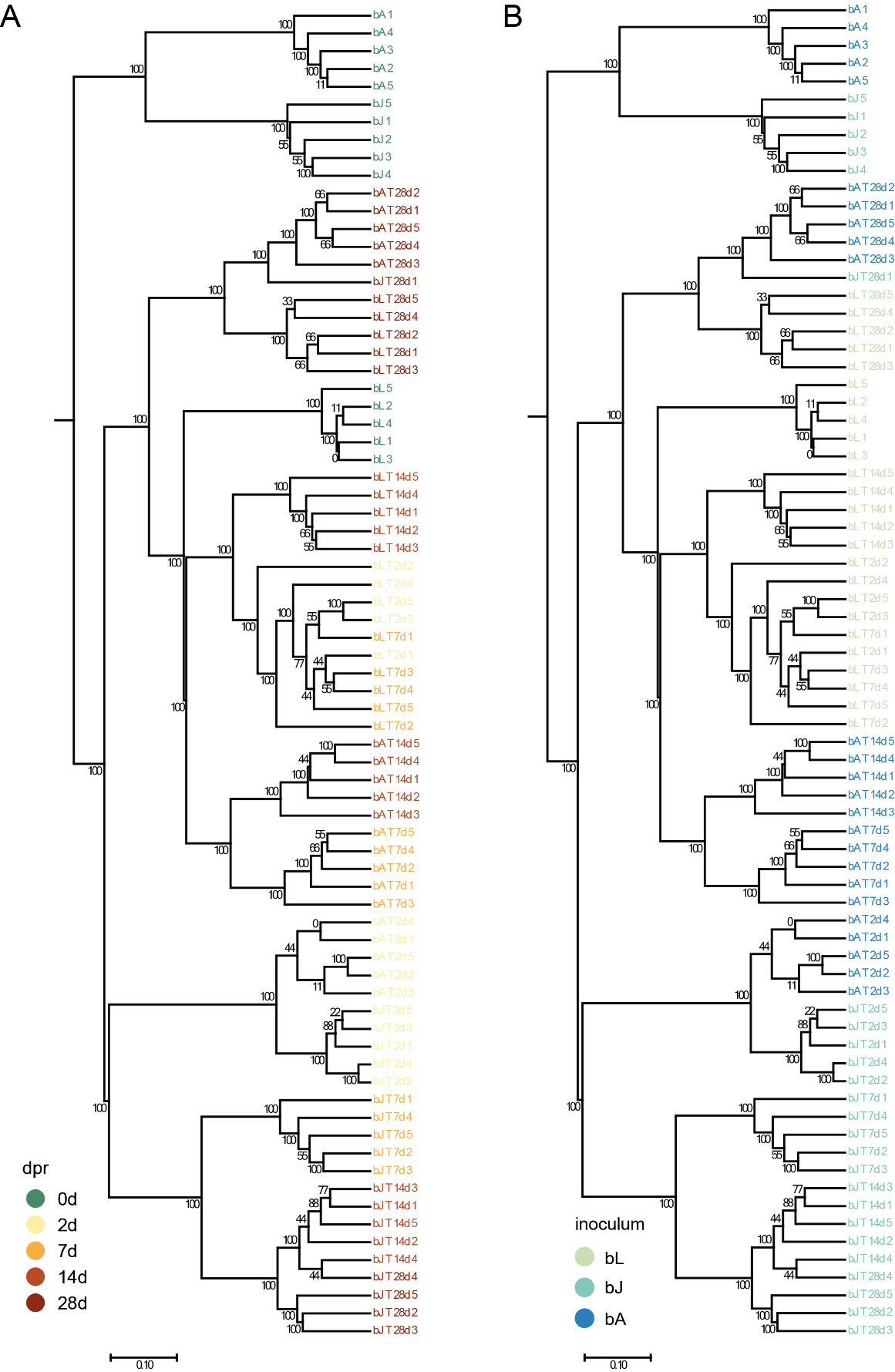

Figure S1: **Analysis of the bacterial recolonization dynamics based on 16S upon recolonization of silicone tubes over the course of one month.** UPGMA tree based on the Bray Curtis dissimilarity for recolonized silicone tubes color-coded according to (A) days post recolonization and (B) inoculum. Samples were taken from the inocula and 2, 7, 14, and 28 days post recolonization.

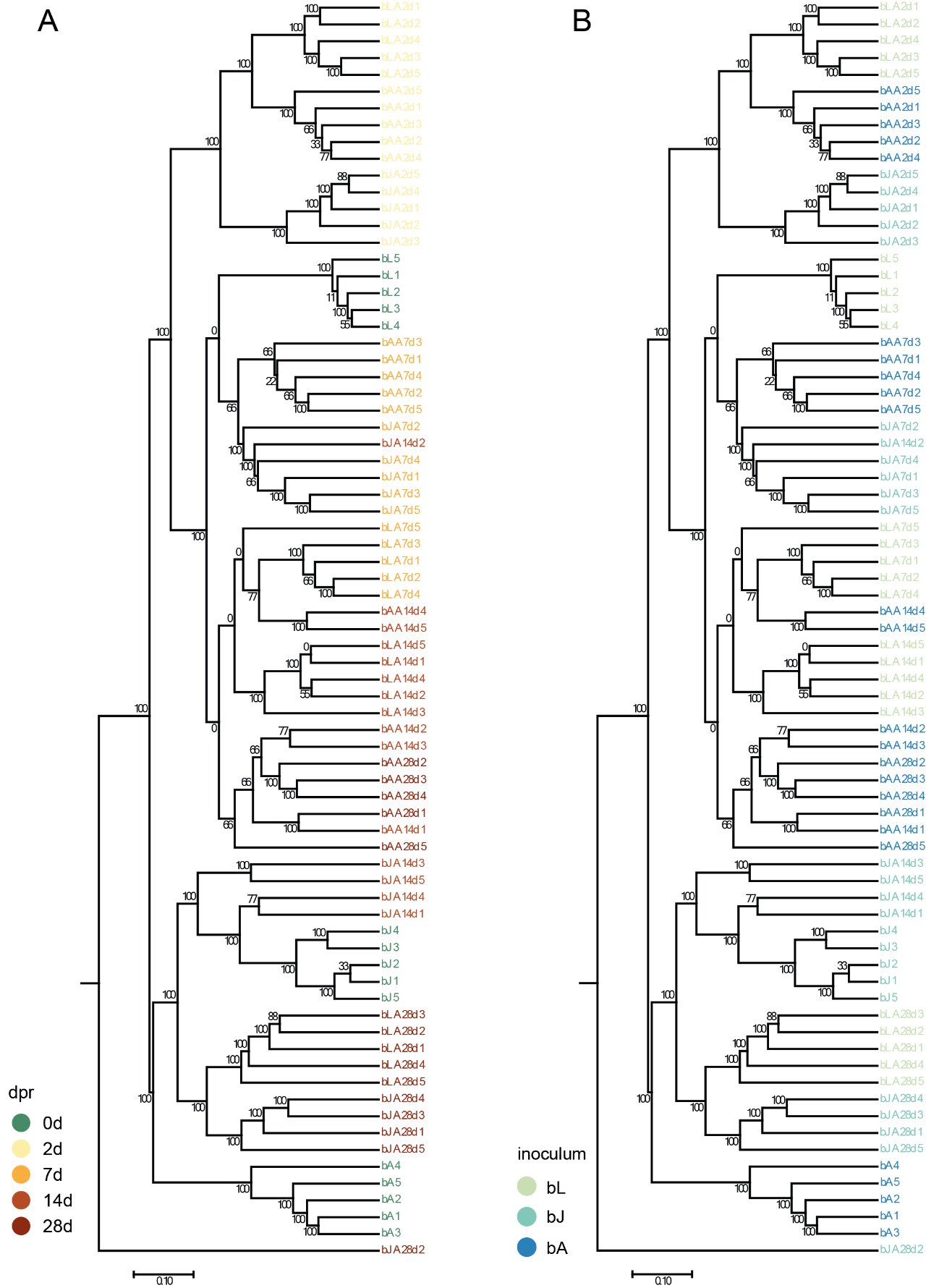

Figure S2: **Analysis of the bacterial recolonization dynamics based on 16S upon recolonization of adult polyps over the course of one month.** UPGMA tree based on the Bray Curtis dissimilarity for recolonized silicone tubes color-coded according to (A) days post recolonization and (B) inoculum. Samples were taken from the inocula and 2, 7, 14, and 28 days post recolonization.

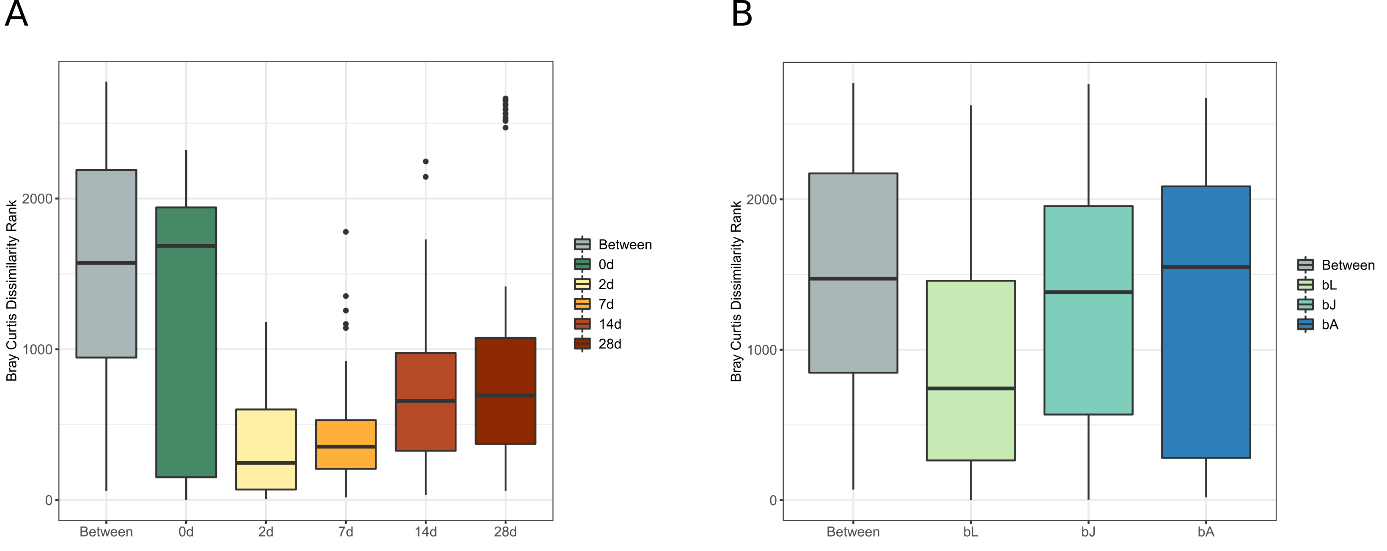

Figure S3: **Bray-Curtis Dissimilarity Ranks of the bacterial community depending on the time and on the inocula.** The ratio of the dissimilarities between and within groups are compared. The compositional dissimilarities between groups are higher than the dissimilarities within the groups, which indicates that the groups are different in their species composition. The higher the R-value, the more dissimilar are the groups. (A) Dissimilarity Ranks according to days post recolonization. Anosim R=0.5804, p=0.001. (B) Dissimilarity Ranks according to inocula. Anosim R=0.2364, p<0.001.

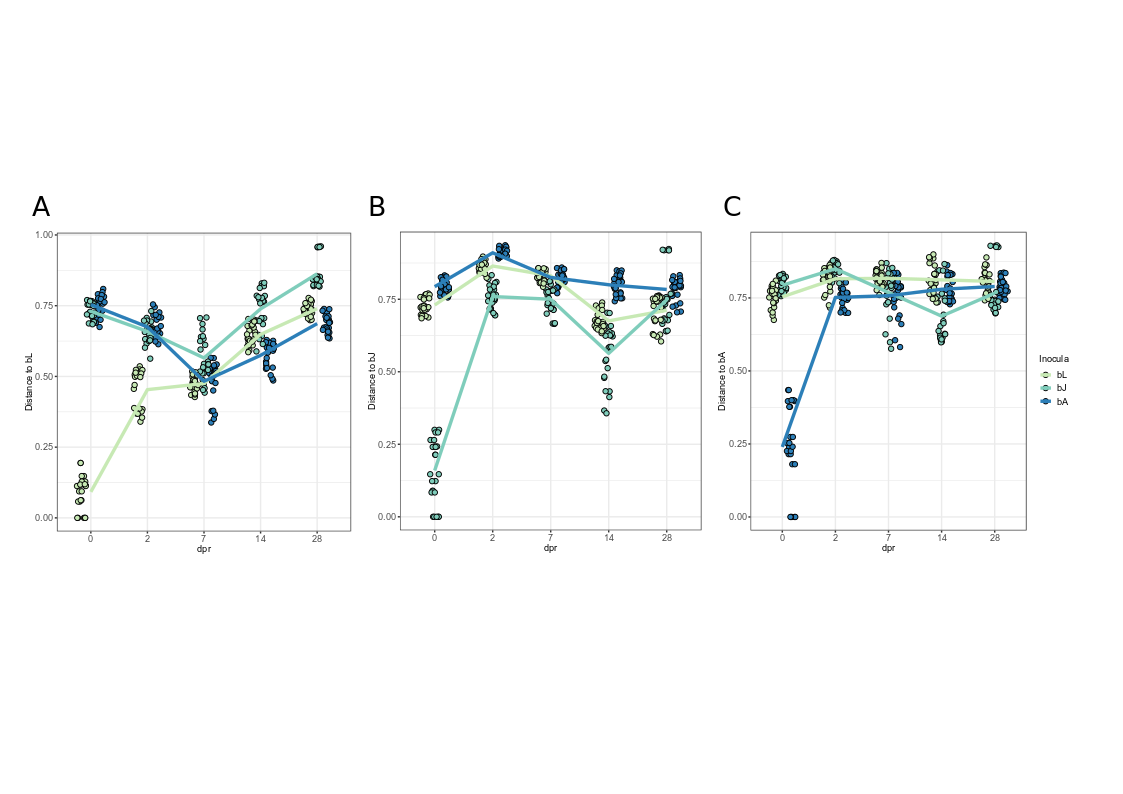

Figure S4: **Bray Curtis distance of the bacterial communities on recolonized animals over time in comparison to the inocula.** Distances are calculated to the three different inocula, respectively. (A) Distance to bL. The lowest distance of all three treatments to the larval inoculum is reached 7 days post recolonization (dpr). (B) Distance to bJ. The lowest distance of all three treatments to the juvenile inoculum is reached 14 dpr. The strongest decline of distance occurred in the polyps recolonized with juvenile bacteria. (C) Distance to bA. There is no decline of distance towards the adult inoculum over the course of this experiment. Solely the polyps recolonized with juvenile bacteria showed a dip in distance 7-14dpr.

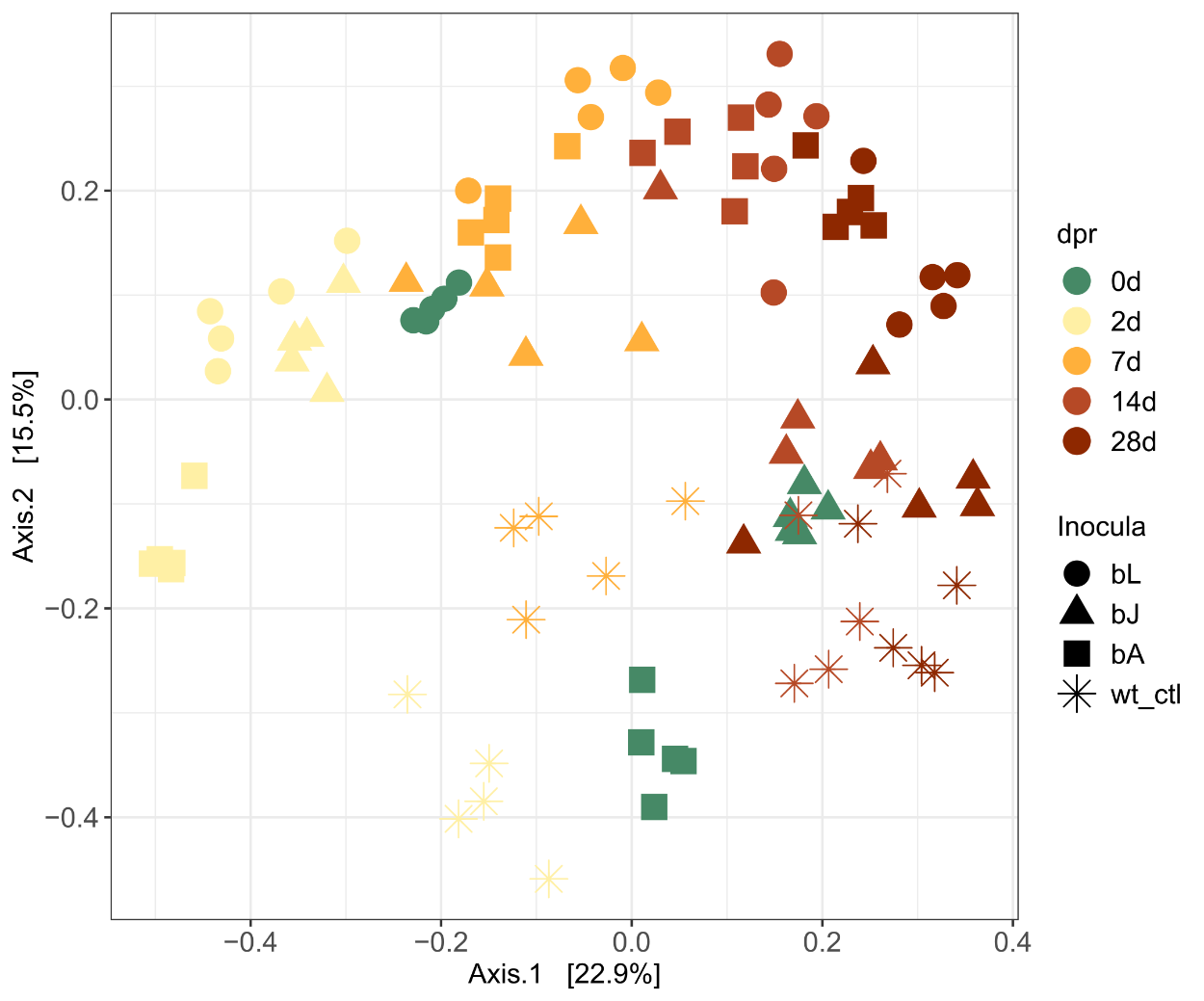

Figure S5: **Recolonization dynamics of gnotobiotic polyps over the course of one month**. In comparison to Figure 1, wildtype samples (wt_ctl) were included in the PCoA plot but dissimilarities are still calculated according to Bray Curtis. Samples of the wildtype control polyps show the effect of time on the microbiome. They move in a circle around the adult inoculum but move towards the juvenile inoculum with time. This effect might pose an effect due to the lack of food.

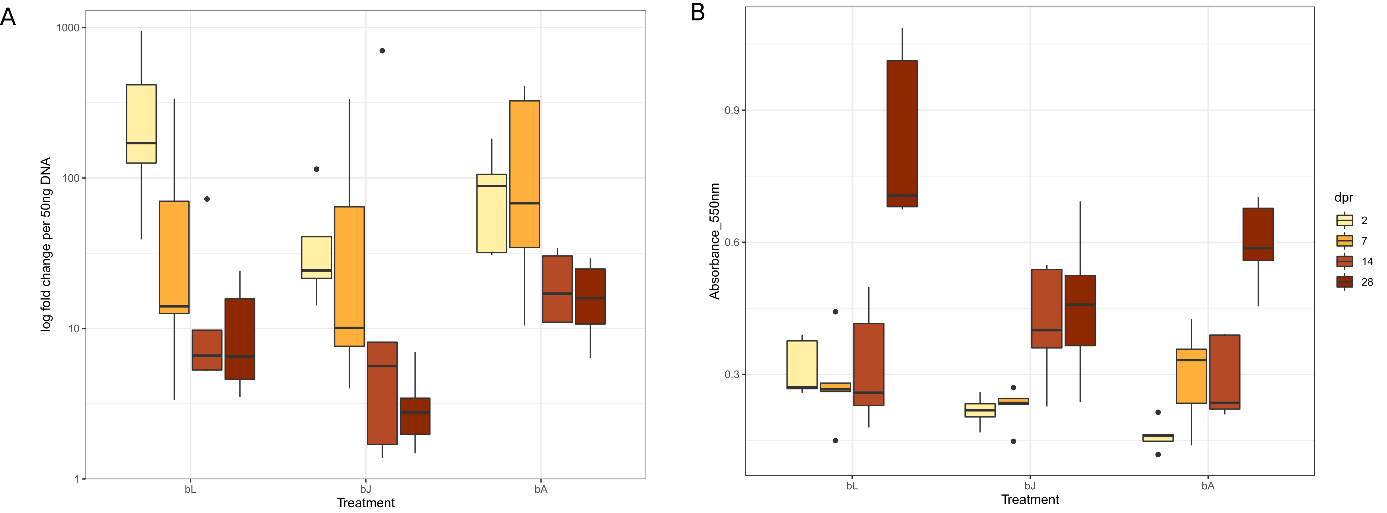

Figure S6: **Absolute bacterial load of the polyps (A) and silicone tubes (B) over the course of the recolonization process**. (A) Bacterial load measured via qRT-PCR on recolonized polyps. The expression of 16S rRNA was normalized to host tissue. Primers for the 16S rRNA gene were specific for the V1-V2 region, Primers for host tissue were specific for the housekeeping gene elongation factor 1alpha. The deltaCT values of all samples were normalized to the values of the wildtype 28dpr. With time, the absolute bacterial load lowers towards the value of the wildtype control animals. (B) Amount of biofilm on silicone tubes quantified by the intensity of crystal violet staining. The intensity was quantified by rising the Crystal Violet off of the silicone tubes and measuring the absorbance at 550 nm. Rise in biofilm formation between 2dpr and 28 dpr was significant in all three treatment groups (p<0.0001). Biofilm formation was just significantly different if bL is compared to bA (p<0.05)

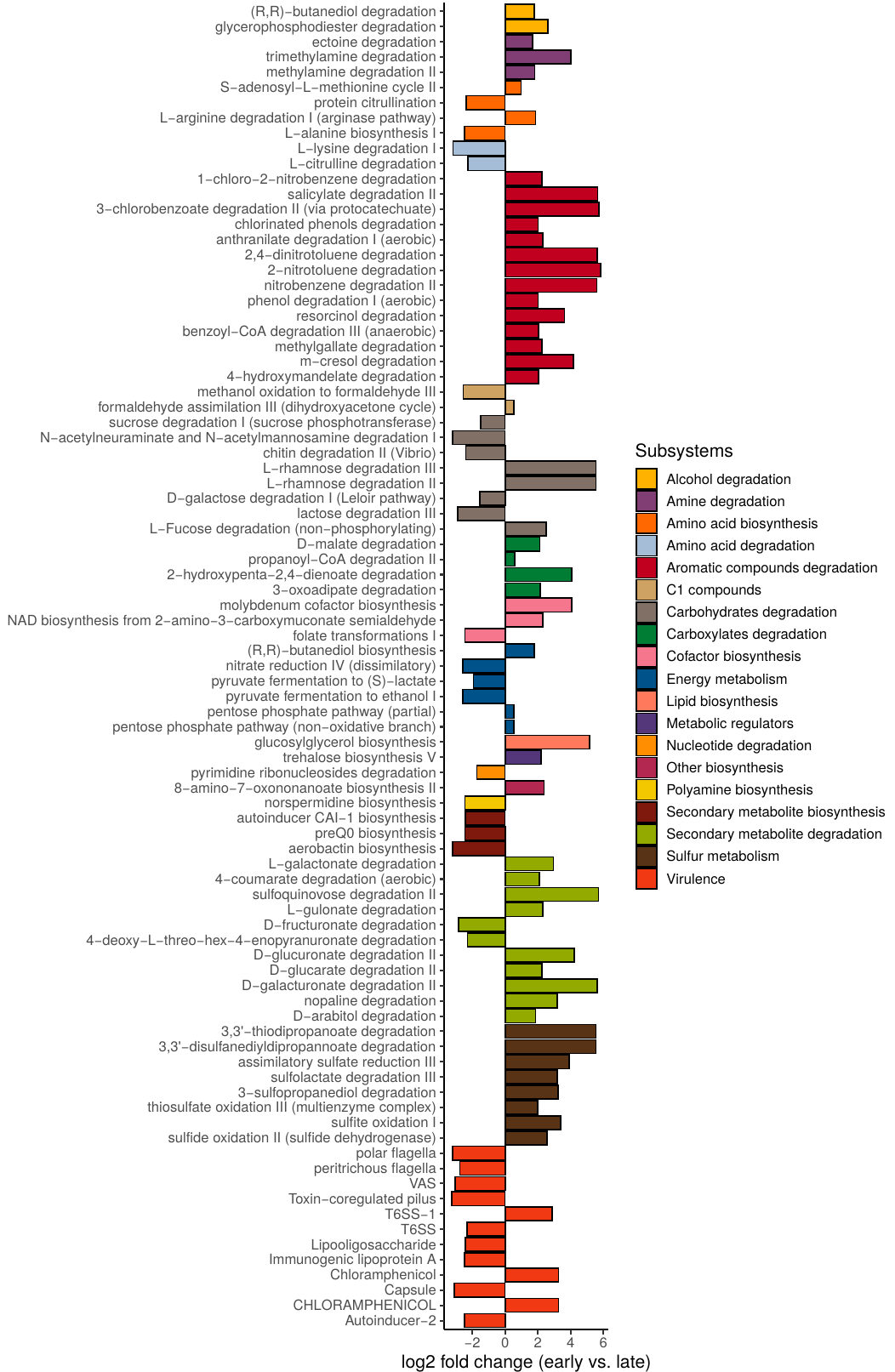

Figure S7: Predicted metabolic and virulence pathway abundances associated stably with early (2d,7d) and late (14d, 28d) colonizer by random forest feature extraction (Boruta). Feature extraction was repeated 100 times and features occurring repeatedly in at least 95% of the cases were considered to be stable. Log2 fold change was calculated from the mean pathway abundances at early time points vs. the mean pathway abundances of later time points.
